# Phase consistency dynamics of memory encoding

**DOI:** 10.1101/2024.12.06.627211

**Authors:** Ryan A. Colyer, Michael J. Kahana

## Abstract

Human and animal studies implicate theta and alpha oscillations in memory function. We tested whether theta, alpha, and beta phase consistency predicts memory encoding dynamics in neurosurgical patients performing delayed free recall tasks with either unrelated (N=188) or categorized words (N=157). We observed widespread post-stimulus phase consistency (3– 21 Hz) and, crucially, identified distinct frequency-specific patterns predictive of successful encoding. Specifically, increased early-list item recall was significantly correlated across subjects with increased theta (3–7 Hz) phase consistency. Subsequent recall analyses, controlling for serial position, revealed distinct frequency signatures for successfully encoded items: theta (3–6 Hz) and alpha (9–14 Hz) for unrelated lists, and theta (3–6 Hz) and beta (14–19 Hz) for categorized lists. Regional analyses for unrelated lists highlighted the lateral temporal cortex for theta effects and the prefrontal cortex for both theta and alpha consistency. These findings provide novel evidence for the frequency-specific presence of increased phase consistency during episodic encoding, revealing its sensitivity to both item context and temporal position within a learning sequence.

**Significance Statement:** Neural oscillations are implicated in human memory encoding, but their precise roles are still being defined. Our study leverages large-scale intracranial EEG datasets from participants undergoing word recall experiments, and introduces analytical innovations for robustly quantifying phase consistency with differing numbers of recalled versus forgotten items. This methodology reveals that phase consistency across different frequency bands (theta, alpha, beta) predicts memory formation. We demonstrate a specific role for theta consistency in encoding early list items and show that the brain adaptively recruits different oscillatory patterns (alpha or beta alongside theta) depending on item context (unrelated vs. categorized lists). These findings advance our understanding of the frequency-specific neural mechanisms supporting human episodic memory, revealing how the brain adapts its encoding strategies based on informational structure.

## Introduction

Numerous studies have highlighted the oscillatory correlates of successful memory encoding across a wide range of frequencies from low theta (3 Hz) through high gamma (100–200 Hz), predominantly via the study of spectral power (Klimesch, 1999; Fell et al., 2001; Sederberg, Kahana, Howard, Donner, & Madsen, 2003; Khader, Jost, Ranganath, & Rösler, 2010; Düzel, Penny, & Burgess, 2010; Headley & Paré, 2017). Furthermore, studies of the causal nature of oscillations in memory function have highlighted the critical nature of phase synchronization within and across brain regions for effective memory function (Hanslmayr, Axmacher, & Inman, 2019). Recent work by ter Wal et al. (2021) has even shown that successful encoding and retrieval for cue-object associative memories corresponded to increased slow theta consistency of phase, and also that encoding and retrieval corresponded to different phases approximately half a cycle apart. This has left the field with little doubt that oscillations correspond to key memory processes, and also shows that the phase synchronizations of these oscillations must be carefully explored to fully elucidate the underlying mechanisms.

The resetting of oscillations following exposure to a stimulus has been observed in both animal (Givens, 1996) and human (Rizzuto et al., 2003) studies during working memory tasks. Rizzuto et al. (2003) presented human participants with a series of temporally jittered consonants in a Sternberg item recognition task, and observed characteristic decreases in power, from a theorized suppression of existing oscillation, correlating with a phase locking identified as the reset of oscillatory phase. These results ranged from 4 Hz through 32 Hz, but were identified as most potent from about 7 Hz through 16 Hz. Rizzuto, Madsen, Bromfield, Schulze-Bonhage, and Kahana (2006) extended this item recognition analysis of reset showing theta phase differences between encoding and retrieval, and attributed this to differences in the phase coding of encoding and retrieval activity (Hasselmo, Bodelon, & Wyble, 2002).

Theoretical debate arose regarding the degree to which components of stimuli- synchronized electrophysiological responses could be attributed to the resetting of phase versus additive responses of traditional event-related potential models, whether this was clearly distinguishable, and how much these were synergistic effects (Sauseng et al., 2007; Telenczuk, Nikulin, & Curio, 2010). To avoid the exclusion of substantial portions of data with filtering approaches attempting to separate these, we take a combined model-independent approach to these two processes, examining the memory-related associates of the consistency of phase taken as a whole, and use this to explore memory dynamics.

The memory processes of encoding are prominently thought to be coordinated throughout the brain by phase synchronization across broad frequency ranges from theta through gamma, with synchronization signals propagating outward from the hippocampus considered a key contributor (Axmacher, Mormann, Fernández, Elger, & Fell, 2006). Oscillatory phase is further modeled as the phenomenon which regulates the timing of information exchange in both local and global networks, with networks at numerous different frequencies operating concurrently (Sauseng & Klimesch, 2008).

Synchronization of neural spike timings with peaks of oscillatory synchronization signals are considered critical to long-term potentiation (Fell & Axmacher, 2011). Schonhaut et al. (2024) recently found, during a spatial navigation task, that neurons in the broader medial temporal lobe phase-lock to hippocampal theta by two complementary processes. One process synchronized to the local field potential, while an additional mechanism was identified that was not mediated by the local field, highlighting the need to carefully explore nuances of phase synchronization. Rutishauser, Ross, Mamelak, and Schuman (2010) also found that the strength of human memory is predicted by the degree of phase-locking between single neuron spike activity and oscillatory phase, confirming the importance of understanding the role of oscillatory phase synchronization in human memory function.

We follow this body of literature by examining the synchronization of phase, measured as phase consistency following stimulus presentation, in a delayed free recall task. A critical factor which influences memory encoding success and cognitive strategies used is the structure of the presented material. Information presented in list form has serial-order effects, such as primacy and recency favoring recall at the begninning and end of the list (Tulving, 2007), which we examined by correlating list-order recall rates with phase consistency across frequencies. Adding semantic structure with categorized lists also has significant impacts on encoding and subsequent recall (Patterson, Meltzer, & Mandler, 1971), which we investigated by comparing serial-position controlled phase consistency effects during the study of lists of unrelated words or semantically categorized words.

## Methods

### Experimental Design

We used data from two delayed free recall tasks conducted with volunteer hospital patients undergoing intracranial electroencephalogram (EEG) monitoring for medication-resistant epilepsy (Kragel et al., 2017; Long et al., 2017; Solomon et al., 2017). Each session consisted of up to 25 lists of 12 words each, presented for 1600 ms with a 0 ms to 250 ms uniform timing jitter added to the first presentation, and a random spacing of 750 ms to 1000 ms between each word. Following this word encoding period, a 20 s self-paced math distractor was provided with problems of summing 3 single digit values, followed by a 30 s free recall period. In the first experiment, uncategorized words were selected from a previously reported wordpool of 300 common English words, with each word presented only once per session (Weidemann et al., 2019). In the second experiment, categorized words were selected from a pool of 300 common words in 25 categories with 12 words per category. Categorized lists were constructed with the category pattern AABBCCBBAACC, with 3 categories used in each list, and each word having one adjacent word of the same category (Weidemann et al., 2019).

We included subjects from our dataset with aggregated sessions for each subject containing a total of 25 or more lists of 12 encoding words each. For our analysis of the whole brain, this resulted in 188 subjects (99 male, 89 female) with 9186 total lists for the uncategorized list experiment, and 157 subjects (88 male, 69 female) with 7195 total lists for the categorized list experiment. We also subdivided electrode contacts by brain region, filtering our subject pool for each analysis to subjects who had contacts available in each region. This yielded an analysis of the hippocampus (Hip.) with 122 subjects comprising 6360 lists for uncategorized, and 111 subjects comprising 5093 lists for categorized. The analysis of the lateral temporal cortex (LTC) had 173 subjects comprising 8424 lists for uncategorized, and 147 subjects comprising 6779 lists for categorized. The third region, the prefrontal cortex (PFC), had 174 subjects comprising 8440 lists for uncategorized, and 143 subjects comprising 6626 lists for categorized. Subjects were able to stop at any point, with common reasons such as fatigue or medical treatment needs, and we included partial sessions in the aggregation of sessions as long as the total count of completed lists met the inclusion criteria.

### Morlet Wavelet Transformation

We analyzed EEG signals from intracranial depth electrodes by using Morlet Wavelet Transformation with the PTSA library. Transformations were performed at integer frequencies from 3 Hz through 24 Hz, from −750 ms prior to word presentation through 750 ms after word presentation. As standard Morlet wavelets which exhibit biases toward 0° and 180° are incompatible with a sensitive phase consistency analysis, the current default PTSA option of zero-integral complete wavelets was used, with a cycle count of 6.

### Phase Consistency

With the aim of better analyzing the properties of phase between measurements of different sample sizes, we developed a new statistical metric for representing the consistency of oscillatory phase. This phase consistency metric was intentionally designed to have the following three properties for a von Mises distribution:

1. A standardized value with a distribution mean-centered around 0 for samples from the uniform distribution, up through a value of 1 for samples with all phase values identical.
2. No systematic bias in metric values due to smaller or larger sample sizes.
3. A valid algebraic mean of metric values such that the average of large numbers of phase consistency metric values approaches the value for the phase distribution from which they were sampled.

This was achieved by first calculating the power-independent square of the average unit-normalized complex plane vectors, *C* and *S*, for each wavelet transformation frequency:

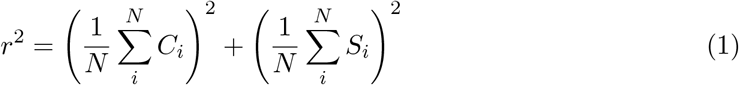

Then we created a phase consistency metric *z*_*s*_ by shifting Rayleigh’s z statistic (Zar, 2010) down by 1, and then scaling down by the number of degrees of freedom given sample size *N* :

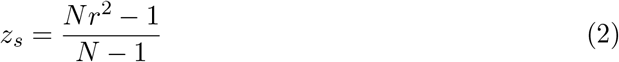

This *z*_*s*_ phase consistency metric provides all three of the above listed properties. These properties were further validated as reliably holding under a series of numerical simulations sampling from the wrappped normal distribution for sample sizes with *N* of 2 and larger.

Note that while nominal phase consistency ranges from 0 to 1, with 0 representing no consistency in phase, and 1 representing perfect consistency in phase, it is an essential property of such a metric that calculated values for individual samples can extend below 0. The most negative value which can be obtained for a sample is − 1, obtainable only with *N* of 2, where the most negative value quickly approaches 0 for larger sample sizes. These negative values represent samples more uniform than the typical sample from a uniform distribution, and are required so that averages sampled from the uniform distribution retain an unbiased central phase consistency of 0.

For electrophysiology data with low signal to noise, phase consistency values trend toward low values, so these statistically clean properties around 0 are essential for using the stated properties in averaging and subtraction operations for combination of data and comparisons between groups.

In Fig. 1 we show simulated sampling from a wrapped normal distribution to highlight visually the amount of bias in phase corresponding to commonly encountered electrophysiological values for phase consistency.

**Figure 1.**
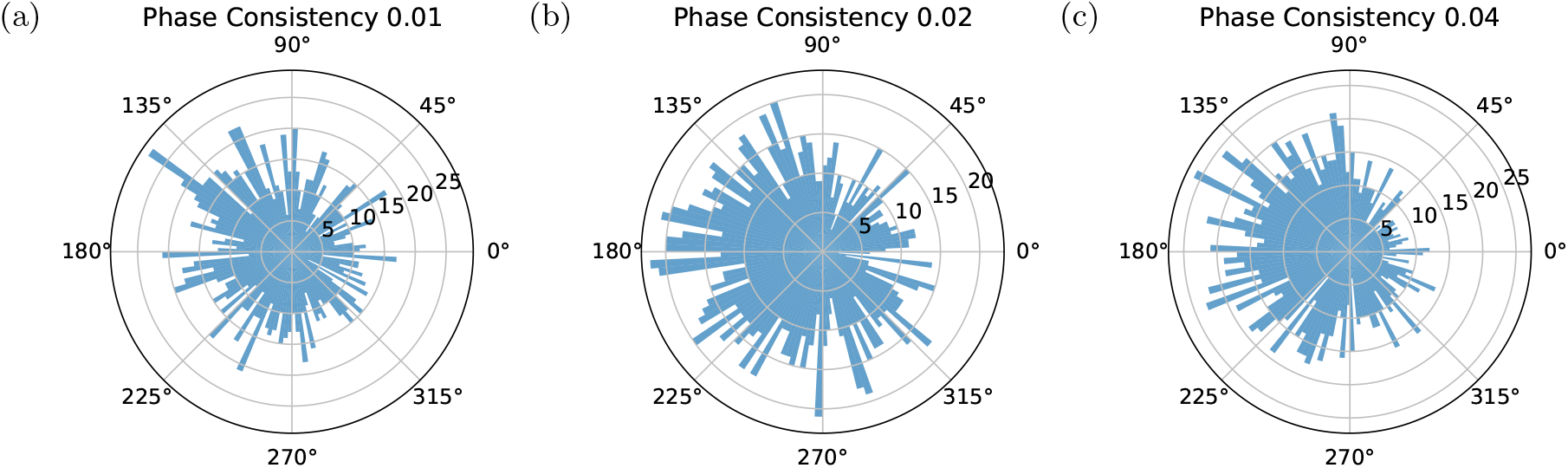
Simulated histograms with N=1600 showing example phase distributions that lead to phase consistency values of **a**. 0.01, **b**. 0.02, and **c**. 0.04.

### Statistical Analysis

Since the means being compared exhibit good normality, significance testing was done as list-count weighted paired t-tests with Benjamini-Hochberg False Discovery Rate (FDR) correction. Confidence bands shown are 95% confidence intervals across subjects.

## Results

Before examining the phase consistency during the encoding of presented words, we examine a baseline comparison in Fig. 2 by looking at the phase consistency at the unjittered start of the countdown which is displayed to participants prior to the word encoding period. Because the start of this event is unjittered, we can see a modest contribution to the consistency of phase appearing at time zero, right at the moment of the countdown start, which indicates a small contribution from participants correctly anticipating when the countdown would begin. The primary effect of phase consistency during this countdown start visual event that participants are not instructed to encode, is a low-frequency focused theta effect centered around a relatively long 375 ms delay after the start of the countdown, with very little higher frequency effect.

**Figure 2.**
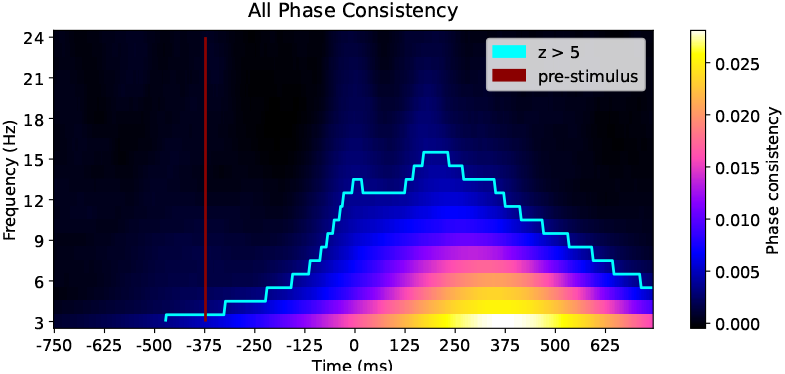
Phase consistency during the unjittered start of the countdown displayed prior to each categorized list presentation.

To begin our main focus, we asked whether words presented during our memory task caused a consistency in the phase of subsequent intracranial EEG signals. We applied the phase consistency metric in Eqn. 2 to time windows from 750 ms before word presentation 750 ms after word presentation, for frequencies from 3 Hz to 24 Hz. The results shown in Fig. 3a and Fig. 3b revealed that phase consistency occurs shortly after stimulus presentation across all the frequencies of consideration. Lower frequencies exhibited a temporally broadened phase consistency result in the analysis due to both the width of those oscillations themselves, and the widths of the Morlet wavelets used to examine them. Due to the jittering of word presentation times built into the task design, phase consistency is driven to an average value of zero prior to the word onset, as the brain cannot synchronize phase to a future event timing that participants cannot predict. The portions of the low frequency response which trail leftward on the plots to before stimulus presentation are due to the broadening from the Morlet wavelets. Figs. 3c–h illustrate phase consistency behavior in three regions: Hippocampus (Hip), Lateral Temporal Cortex (LTC), and Prefrontal Cortex (PFC).

**Figure 3.**
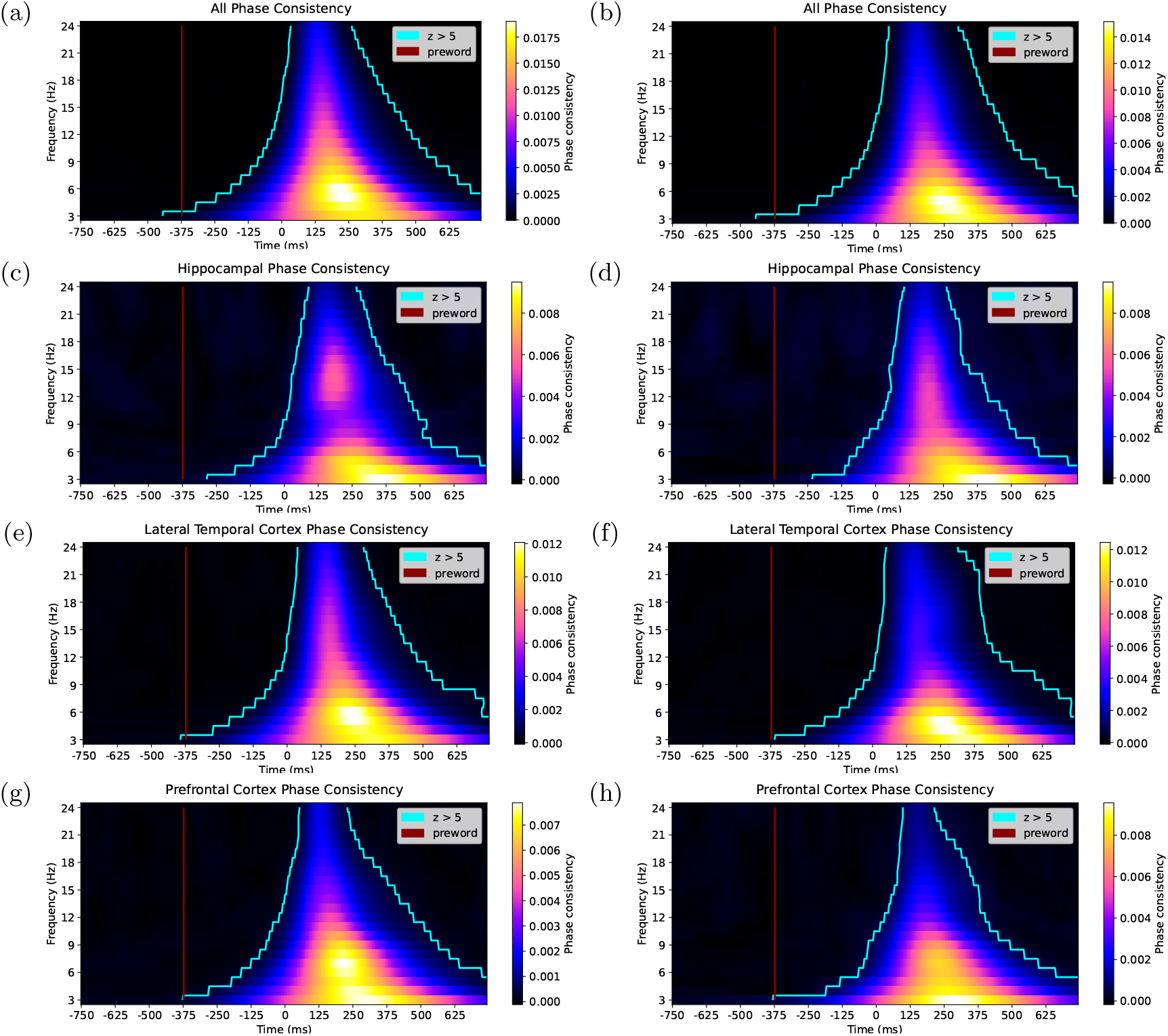
Average phase consistency across events and all subjects for **(a**,**b)** all electrode contacts, **(c**,**d)** hippocampal contacts, **(e**,**f)** LTC contacts, and **(g**,**h)** PFC contacts, for **(a**,**c**,**e**,**g)** uncategorized lists and **(b**,**d**,**f**,**h)** categorized lists.

Having established that phase consistency appears broadly throughout the brain following item presentation, we next asked whether the degree of phase consistency would predict aspects of memory for the studied items. As the serial position of a studied item strongly modulates subsequent memory, we first examined phase consistency as a function of serial position. We then consider whether — controlling for serial position — phase consistency differs between the encoding of subsequently remembered and forgotten items.

Fig. 4a and Fig. 4d illustrate phase consistency both before and after item presentation as a function of serial position. Whereas the pre-word phase consistency reliably overlaps zero, the post-word phase consistency is strongly positive, and follows a pattern similar to the serial position curve of recall rates.

**Figure 4.**
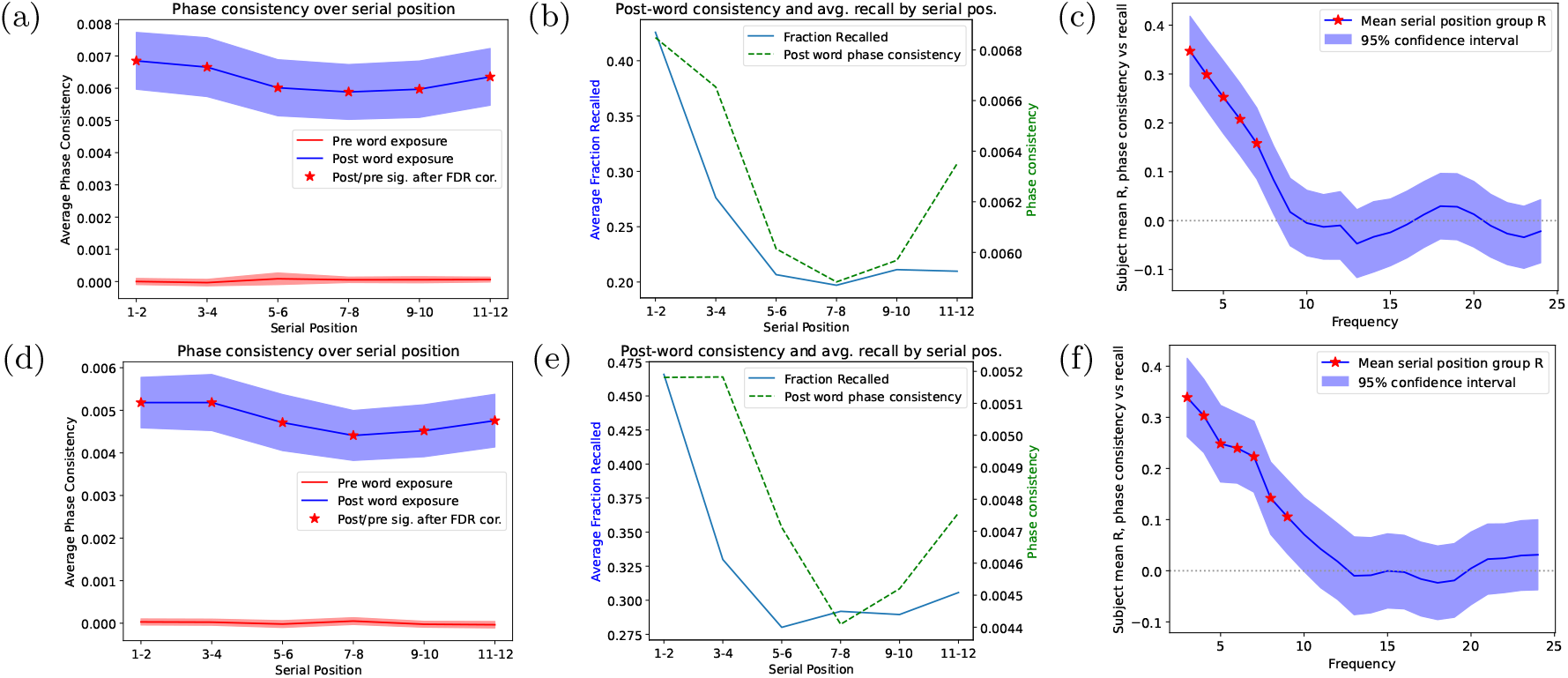
**(a**,**d)** Whole brain phase consistency with 95% confidence intervals averaged over all frequencies from the pre-word time window of −750 ms to −375 ms compared to the post-word time window of 125 ms to 500 ms for each serial position grouping. **(b**,**e)** Post-word phase consistency for each serial position grouping overlayed with average fraction correct for each grouping. **(c**,**f)** Mean across subjects of the correlation R between serial position group recall rates and the post-pre phase consistency difference at each frequency, with 95% confidence intervals and significance marked after FDR correction. For **(a**,**b**,**c)** uncategorized and **(d**,**e**,**f)** categorized lists.

For Fig. 4b and 4e we plotted both the fraction recalled and the phase consistency vertically aligned with their respective maximum and minimum values, which shows visually the relationship over all subjects between the average post-pre phase consistency difference and the average recall rate across the serial position groups. We assessed the reliability of this relationship across subjects by performing correlations for each subject between the recall rate of the serial position groups and the phase consistency difference between the post-word and pre-word regions over all frequencies from 3–24 Hz. Fig. 5 illustrates the distributions of these correlation values and evaluates whether the mean correlation (across subjects) reliably differs from zero. The 95% confidence intervals shown for the mean correlation are small due to the sum over 188 and 157 subjects for the two datasets, complying with a rough approximation of the expected interval width from 2 over the square root of subject counts. For the entire brain we see a reliably positive correlation between phase consistency and serial position – an effect that appears separately in each of the three regions of interest (hippocampus, temporal cortex and prefrontal cortex). Thus, as recall rates appear highest for early list positions we also find greater stimulus-related phase consistency for early list positions.

**Figure 5.**
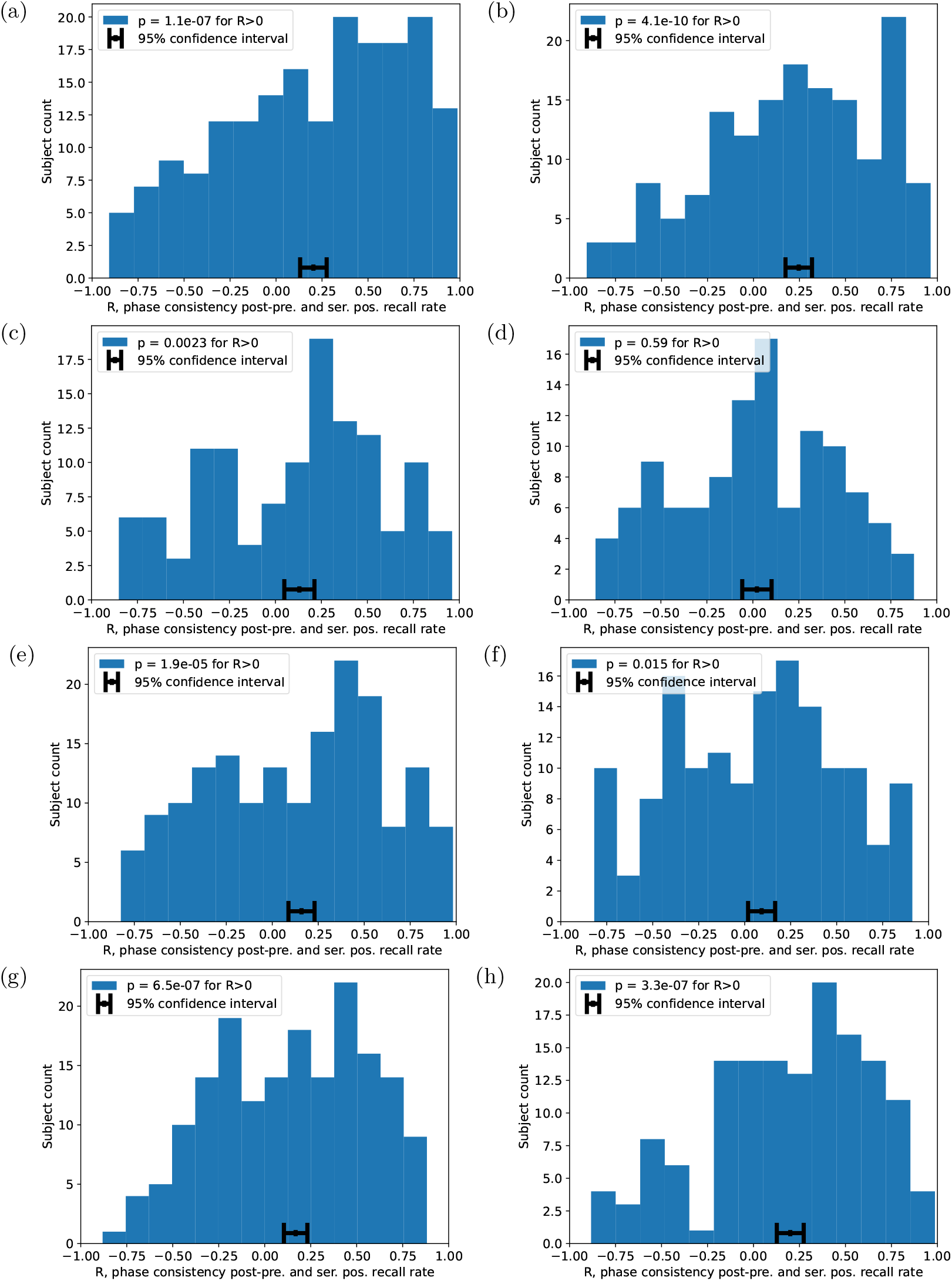
Histograms of subject correlation values between the successful recall fraction for serial position groups of two, and the phase consistency difference between post-word and pre-word regions. Correlations were significantly positive across subjects for each of **(a**,**b)** whole brain, **(c**,**d)** Hip., **(e**,**f)** LTC, **(g**,**h)** and PFC, for **(a**,**c**,**e**,**g)** uncategorized and **(b**,**d**,**f**,**h)** categorized lists.

We then computed serial position grouping correlations between recall rates and post-pre phase consistency differences for each frequency separately, finding the mean correlation value for each frequency, and produced the frequency dependence of this serial position effect shown in Fig. 4c. This plot reveals that the serial position phase consistency effect exists primarily within the band of 3 Hz through 7 Hz, and convincingly overlaps zero at higher frequencies. In additional data not shown, this effect follows an identical pattern across the subdivided brain regions of Hip., LTC, and PFC, although with the significance of the matching trend not surviving FDR correction within the sparser hippocampal data. This convincingly reveals that across the brain, the phase consistency correlates of serial position effects on subsequent recall rates are within the theta bands of 3–7 Hz, with the strongest effects at the lower frequencies.

Given that serial position strongly influences recall probability, we chose to first establish the nature of serial position effects of phase consistency, and then control for serial position to rule out the possibility that the serial position correlation is the exclusive driver of phase consistency effects. Fig. 6a shows the frequency distribution of the overall subsequent memory effect (SME) across the whole brain, without controlling for serial position. It can be seen that all frequencies examined contribute to the overall effect across all conditions, with sensitivity to observing phase consistency trailing off at the highest frequencies above 21 Hz. By separately calculating the phase consistency SME differences in the primacy, recency, and middle regions, we are able to address the biases that would be created in the SME by the higher recall rates in primacy and recency portions creating a higher weighting of those parts of the serial position curve in the recalled data than in the non-recalled data. In Fig. 6b we show the SME controlled for serial position effects by grouping early list, middle list, and late list items separately, and then merging the resulting phase consistency differences between recalled and not-recalled items. This result gives high confidence in an overall remaining SME across the whole brain and all frequencies even after controlling for serial position effects.

**Figure 6.**
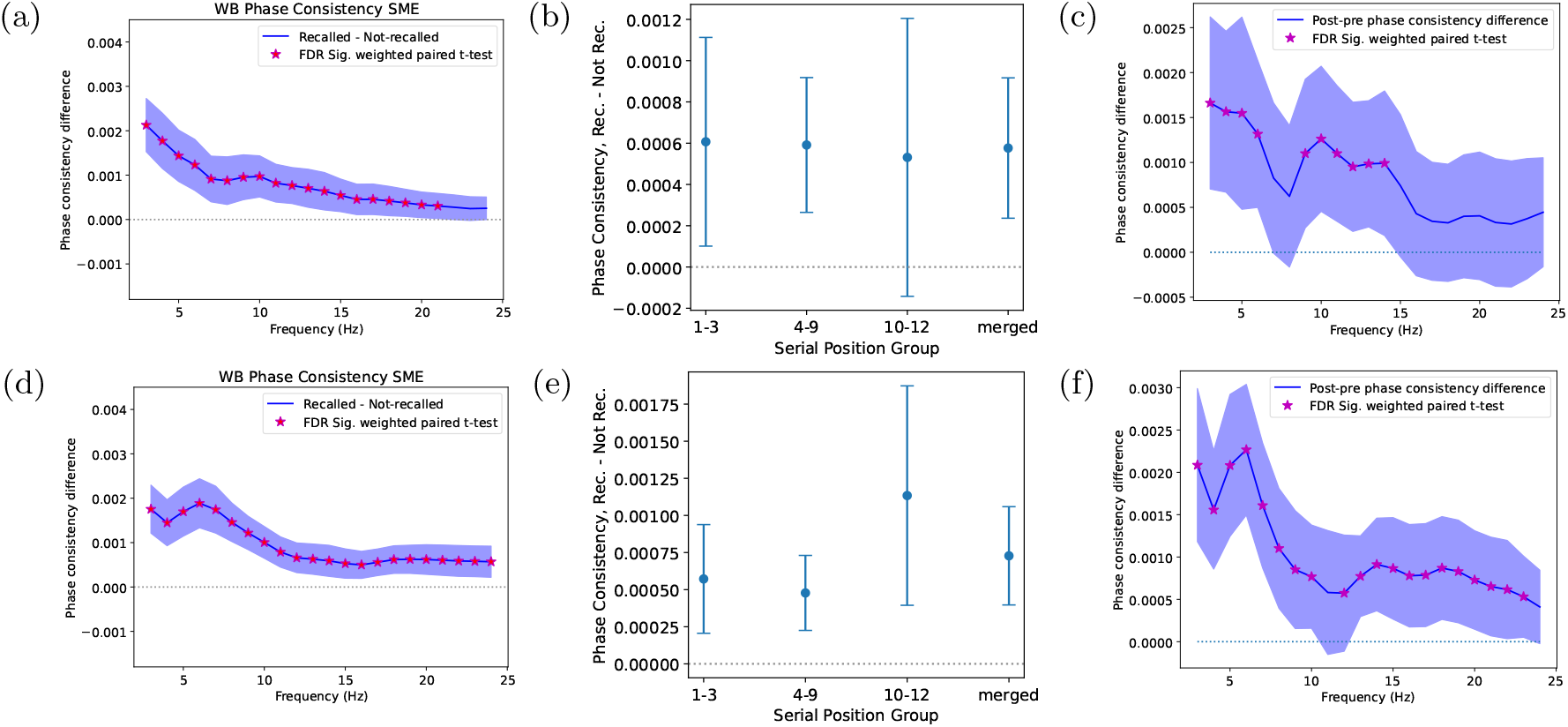
Whole brain phase consistency SME (recalled minus not-recalled) with 95% confidence intervals **(a**,**d)** as a frequency distribution without serial position control, **(b**,**e)** overall effect controlling for serial position grouping, and **(c**,**f)** as a frequency distribution controlling for serial position grouping. For **(a**,**b**,**c)** uncategorized and **(d**,**e**,**f)** categorized lists.

We then performed this serial position control analysis for each frequency separately, resulting in Fig. 6c, where a significant serial position controlled SME appears across both theta (3–6 Hz) and alpha (9–14 Hz) bands. This pronounced alpha component of the phase consistency SME is distinct from the theta-only effect of the serial position correlate found in 4c.

Fig. 7a, 7c, and 7e show the detailed whole brain SME for phase consistency separated by frequency and serial position grouping, where early list items exhibit a theta SME from 3–6 Hz, middle list items include that band plus an alpha effect at 9–10 Hz and a beta effect at 16–20 Hz, and late list items have an alpha SME from 9–11 Hz. Given the serial position controlled SME effect of Fig. 6c highlighting the overall significance of the theta and alpha phase consistency components, this serial position division of the data shows the theta phase consistency SME appearing in the earlier list items and the alpha phase consistency SME appearing in the later list items.

**Figure 7.**
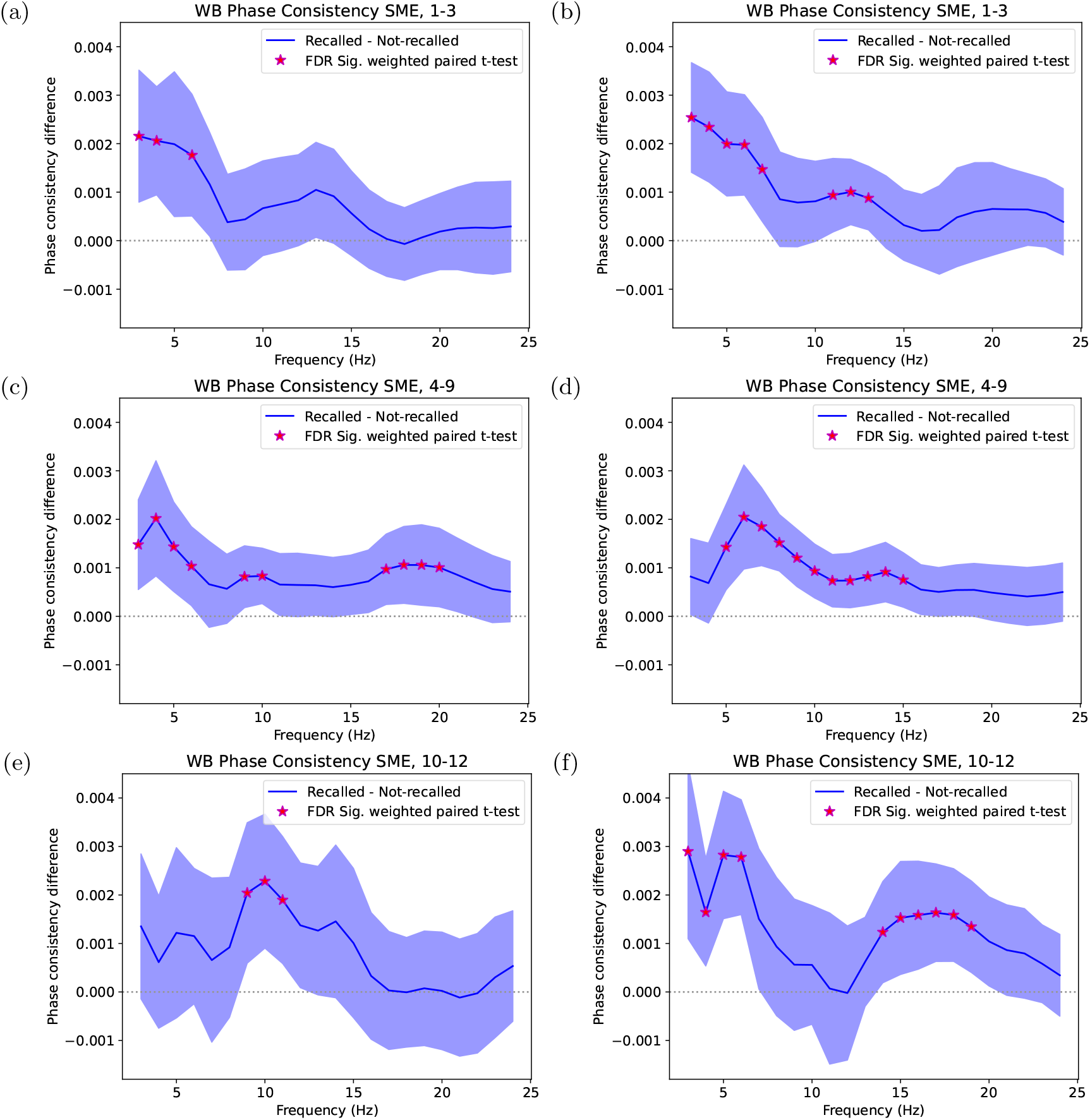
Phase consistency SME by serial position grouping from recalled minus not-recalled, averaged over all contacts and all subjects. Serial positions **(a**,**b)** 1–3, **(c**,**d)** 4–9, and **(e**,**f)** 10–12, for **(a**,**c**,**e)** uncategorized and **(b**,**d**,**f)** categorized lists.

We also separated the SME across the other dimension of brain region without controlling for serial position. Fig. 8a and Fig. 8b show the hippocampal contacts trending upward in their SME at lower theta frequencies as one might expect, but without significance due to the smaller number of total contacts contributing to this result. Fig. 8c and Fig. 8d shows only a significant theta phase consistency SME in the LTC, while in Fig. 8e for uncategorized lists there is a bimodal theta phase consistency (3–4 Hz) and an extended alpha phase consistency (6–11 Hz) SME.

**Figure 8.**
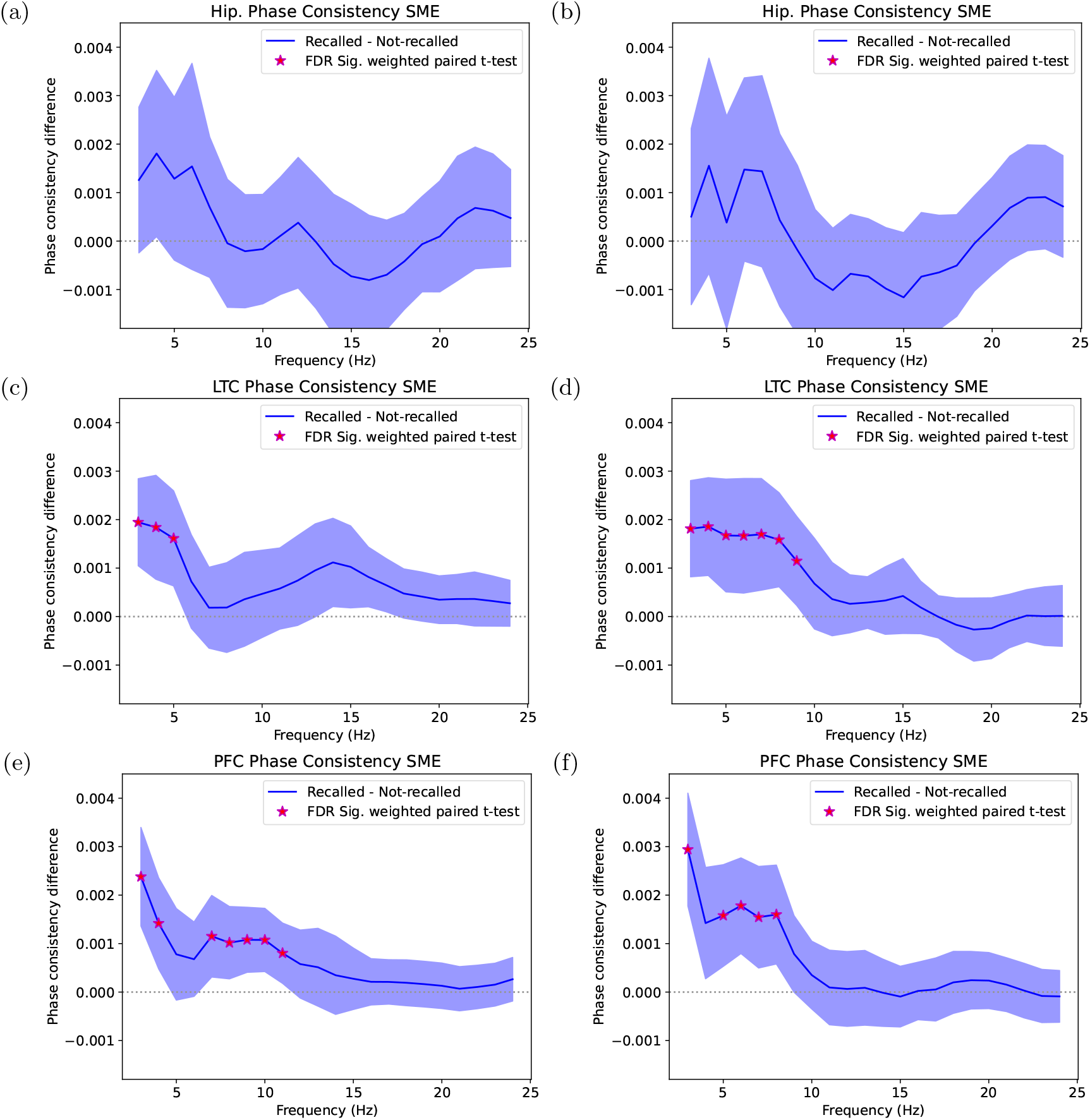
Phase consistency SME by serial position grouping from recalled minus not-recalled, averaged over all contacts and all subjects. **(a**,**b)** Hippocampus, **(c**,**d)** LTC, and **(e**,**f)** PFC, for **(a**,**c**,**e)** uncategorized and **(b**,**d**,**f)** categorized lists.

## Discussion

We showed that theta phase consistency rises significantly from 3–7 Hz in correlation with serial position effects primarily driven by primacy in a delayed free recall task. Furthermore we showed that after controlling for serial position, a significant subsequent memory effect of theta phase consistency increase (3–6 Hz) combined with alpha phase consistency increase (9–14 Hz) for uncategorized lists, and primarily theta and beta (14–19 Hz) for categorized lists. The observation that serial position correlates are confined to lower frequencies, while SME effects after controlling for serial position for uncategorized lists include alpha, and for categorized lists include higher frequencies strongest in the beta bands, shows that there are differing mechanisms of oscillatory phase consistency associated with the increased recall of primacy, versus increases in subsequent recall when controlling for serial position.

Because word presentations were jittered in their timing, we were able to conclude these results are causally due to participant responses to the word presentation events. Confirming that phase consistency values prior to the word presentation have confidence bands that strongly overlap zero and are well separated from the other effects validates the expectation that the remaining effects are specifically due to participant responses to the word presentations, as was observed clearly in Fig. 4a. We propose that this strong theoretical guarantee, with a built-in validation, provides a useful foundation for exploring oscillatory responses.

With the above results we have demonstrated the general potency of the phase consistency metric in Eqn. 2 for studying oscillatory phase relationships, and encourage its broader use. This approach is not limited to studying temporal synchronization with stimuli, but equivalently provides a convenient tool for examining the consistency of relative phase, such as between contacts.

We have shown in a free recall task that oscillatory phase consistency 125 ms to 500 ms following word presentation appears throughout the brain across a broad frequency range from 3 Hz through 21 Hz. The strongest magnitudes of measured phase consistency exhibited in Fig. 3, without consideration of any memory dynamics, were found in the theta frequency range. However, interpreting the relative magnitudes of far away frequencies of phase consistency must consider the greater ease of extracting phase consistency at lower frequencies. This improved ease of extracting consistency is facilitated both by longer low-frequency Morlet wavelets having longer time periods to integrate data over, as well as a reduced sensitivity of lower frequency consistency measurements to small fluctuations in experimental synchronization or physiological fluctuations in response timing. Griffiths and Jensen (2023) recently reviewed the episodic memory role of synchronization of gamma oscillatory activity (30–80 Hz) in conjunction with theta, and it is notable that directly examining the consistency of phase with respect to a stimulus in that higher frequency band is much less sensitive due to the higher sensitivity to intertrial differences in response times. We were unable to observe meaningful phase consistency above the 24 Hz range shown, which we attributed to faster frequencies having greater sensitivity to physiological response timing fluctuations. However we expect this method would extend better to higher frequencies if rather than observing synchronization of phase with respect to external stimulus timing, the same method were applied to the synchronization of relative phase between contacts, which would control for physiological response timing fluctuations. We also propose that using this consistency metric for contact pairings would enable longer term tracking of the temporal dynamics of responses to an event.

We demonstrated a particularly clear correspondence between theta phase consistency (3–7 Hz) and the recall rates at various serial positions. There is a strong theoretical reason to have expected this correspondence for list primacy. Hasselmo et al. (2002) suggests that increased theta phase consistency corresponding to subsequently remembered items, along with increased theta phase consistency corresponding to attention, providing support for an attentional explanation for the higher subsequent recall. ter Wal et al. (2021) also suggests that both visual attention and mnemonic attention are coordinated by a reset in theta oscillations. As attentional effects are expected to contribute to increased recall rates during primacy portions of a list, our finding that the serial position effect is focused on consistency of theta lends support to these explanations.

The apparent correspondence in Fig. 4b and Fig. 4e of phase consistency with the slightly increased recall rate in late list items in a delayed free recall task requires a more nuanced consideration, but is plausibly due to participant encoding behaviors or strategies that arise upon the expectation of approaching the end of the consistently sized lists. In Fig. 8e, the phase consistency SME for this late list portion of the data showed a characteristically different frequency profile for the contribution to subsequent recalls than was observed for the primacy regions, with significance in the alpha band that was found to correspond to the SME. This did not appear in the frequency plot of Fig. 4c, which showed only a theta effect, as due to the math distractor task in the delayed recall experiment causing the serial position curve to be dominated by primacy, the correlation examined was expected to be dominated by the much stronger primacy effect on recall probability.

Even after controlling for serial position, phase consistency showed a clear theta and alpha phase consistency SME, distinct from the primacy effect, supporting the notion that phase consistency plays a multifaceted role in the processes that synchronize the brain during the task of encoding word presentation. The regional division of the data showed a theta effect in the LTC, and a dual theta/alpha effect in the PFC for uncategorized lists. A more nuanced analysis of the regional contributions to these processes was hindered by variability and reduced data counts after restricting to electrode contacts within specific brain regions. Additional analyses not shown, attempting to divide across all the axes of frequency, brain region, and serial position grouping, were washed out by the sparsity of data, but might yield interesting insights with additional data or alternative analytical approaches or modeling.

A particularly encouraging aspect of these phase consistency analyses was that the effects observed were both potent and widespread, indicating the potential of using this information as an independent data axis to oscillatory power for combined biomarkers of successful encoding in machine learning approaches.

Overall, this phase consistency approach to exploring phase consistency in a delayed free recall task revealed that phase consistency corresponds to successful recall in at least two different ways, a widespread theta phase consistency effect associated with the primacy effect on recall, and dual theta/alpha (semantically unrelated) and theta/beta (semantically categorized) phase consistency effects independent of serial order exhibiting portions of these components in the LTC and PFC. These results show that there are multiple mechanisms by which phase consistency associates with the memory dynamics leading to the successful recall of episodic associations, and warrant further study to explore the memory networks related to this synchronization.

## Data and Code Accessibility

All experimental data is on OpenNeuro at https://openneuro.org/datasets/ds004789 and https://openneuro.org/datasets/ds004809, and our analysis code can be freely obtained at: https://github.com/rcolyer/PhaseConsistency2025

## Acknowledgments

We are grateful to the patients for their generous participation in this project, and wish to thank the numerous hospital staff and researchers who were involved in data acquisition. This work was supported by the NIH grant MH55687.

## References

Axmacher, N., Mormann, F., Fernández, G., Elger, C. E., & Fell, J. (2006). Memory formation by neuronal synchronization. Brain Research Reviews, 52 (1). doi: 10.1016/j.brainresrev.2006.01.007

Düzel, E., Penny, W. D., & Burgess, N. (2010). Brain oscillations and memory. Current Opinion in Neurobiology, 20 (2), 143–149.

Fell, J., & Axmacher, N. (2011, February). The role of phase synchronization in memory processes. Nature Reviews. Neuroscience, 12 (2), 105–118.

Fell, J., Klaver, P., Lehnertz, K., Grunwald, T., Schaller, C., Elger, C. E., & Fernandez, G. (2001). Human memory formation is accompanied by rhinal-hippocampal coupling and decoupling. Nature Neuroscience, 4 (12), 1259–1264. doi: 10.1038/nn759

Givens, B. (1996). Stimulus-evoked resetting of the dentate theta rhythm: relation to working memory. Neuroreport, 8, 159–163.

Griffiths, B. J., & Jensen, O. (2023). Gamma oscillations and episodic memory. Trends in Neurosciences.

Hanslmayr, S., Axmacher, N., & Inman, C. S. (2019). Modulating human memory via entrainment of brain oscillations. Trends in Neurosciences, 42 (7), 385–499. doi: 10.1016/j.tins.2019.04.004

Hasselmo, M. E., Bodelon, C., & Wyble, B. P. (2002). A proposed function for hippocampal theta rhythm: Separate phases of encoding and retrieval enhance reversal of prior learning., 14, 793–817.

Headley, D. B., & Paré, D. (2017). Common oscillatory mechanisms across multiple memory systems. npj Science of Learning, 2 (1), 1.

Khader, P. H., Jost, K., Ranganath, C., & Rösler, F. (2010). Theta and alpha oscillations during working-memory maintenance predict successful long-term memory encoding. Neuroscience letters, 468 (3), 339–343.

Klimesch, W. (1999). EEG alpha and theta oscillations reflect cognitive and memory performance: a review and analysis. Brain Research Reviews, 29, 169–195.

Kragel, J. E., Ezzyat, Y., Sperling, M. R., Gorniak, R., Worrell, G. A., Berry, B. M., … Kahana, M. J. (2017). Similar patterns of neural activity predict memory function during encoding and retrieval. NeuroImage, 155, 60–71. doi: 10.1016/j.neuroimage.2017.03.042

Long, N. M., Sperling, M. R., Worrell, G. A., Davis, K. A., Gross, R. E., Lega, B. C., … Kahana, M. J. (2017). Contextually mediated spontaneous retrieval is specific to the hippocampus. Current Biology, 27 (7), 1074–1079. doi: 10.1016/j.cub.2017.02.054

Patterson, K. E., Meltzer, R. H., & Mandler, G. (1971). Inter-response times in categorized free recall. Journal of Verbal Learning and Verbal Behavior, 10, 417–426. doi: 10.1016/S0022-5371(71)80041-5

Rizzuto, D., Madsen, J. R., Bromfield, E. B., Schulze-Bonhage, A., & Kahana, M. J. (2006). Human neocortical oscillations exhibit theta phase differences between encoding and retrieval. NeuroImage, 31 (3), 1352–1358.

Rizzuto, D., Madsen, J. R., Bromfield, E. B., Schulze-Bonhage, A., Seelig, D., Aschenbrenner-Scheibe, R., & Kahana, M. J. (2003). Reset of human neocortical oscillations during a working memory task. Proceedings of the National Academy of Sciences, USA, 100 (13), 7931–7936.

Rutishauser, U., Ross, I., Mamelak, A., & Schuman, E. (2010). Human memory strength is predicted by theta-frequency phase-locking of single neurons. Nature, 464 (7290), 903–907.

Sauseng, P., & Klimesch, W. (2008). What does phase information of oscillatory brain activity tell us about cognitive processes? Neuroscience & Biobehavioral Reviews, 32 (5), 1001–1013.

Sauseng, P., Klimesch, W., Gruber, W. R., Hanslmayr, S., Freunberger, R., & Doppelmayr, M. (2007). Are event-related potential components generated by phase resetting of brain oscillations? a critical discussion. Neuroscience, 146 (4), 1435–1444.

Schonhaut, D. R., Rao, A. M., Ramayya, A. G., Solomon, E. A., Herweg, N. A., Fried, I., & Kahana, M. J. (2024). MTL neurons phase-lock to human hippocampal theta. eLife, 13, e85753. doi: 10.1101/2020.06.30.180174

Sederberg, P. B., Kahana, M. J., Howard, M. W., Donner, E. J., & Madsen, J. R. (2003). Theta and gamma oscillations during encoding predict subsequent recall. Journal of Neuroscience, 23 (34), 10809–10814. doi: 10.1523/JNEUROSCI.23-34-10809.2003

Solomon, E. A., Kragel, J. E., Sperling, M. R., Sharan, A. D., Worrell, G. A., Kucewicz, M. T., … Kahana, M. J. (2017). Widespread theta synchrony and high-frequency desynchronization underlies enhanced cognition. Nature Communications, 8 (1), 1704. doi: 10.1038/s41467-017-01763-2

Telenczuk, B., Nikulin, V. V., & Curio, G. (2010). Role of neuronal synchrony in the generation of evoked eeg/meg responses. Journal of neurophysiology, 104 (6), 3557– 3567.

ter Wal, M., Linde-Domingo, J., Lifanov, J., Roux, F., Kolibius, L. D., Gollwitzer, S., … others (2021). Theta rhythmicity governs human behavior and hippocampal signals during memory-dependent tasks. Nature Communications, 12 (1), 7048.

Tulving, E. (2007). Memory and mind: A festschrift for gordon h. bower. In M. Gluck, J. Anderson, & S. M. Kosslyn (Eds.), (chap. On the law of primacy). New Jersey: Lawrence 20 Erlbaum Associates.

Weidemann, C. T., Kragel, J. E., Lega, B. C., Worrell, G. A., Sperling, M. R., Sharan, A. D., … Kahana, M. J. (2019). Neural activity reveals interactions between episodic and semantic memory systems during retrieval. Journal of Experimental Psychology: General, 148 (1), 1–12. doi: 10.1037/xge0000480

Zar, J. H. (2010). Biostatistical analysis, fifth edition. New Jersey: Practice Hall.

